# Autism is associated with inter-individual variations of gray and white matter morphology

**DOI:** 10.1101/2022.02.16.480649

**Authors:** Ting Mei, Natalie J. Forde, Dorothea L. Floris, Flavio Dell’Acqua, Richard Stones, Iva Ilioska, Sarah Durston, Carolin Moessnang, Tobias Banaschewski, Rosemary J. Holt, Simon Baron-Cohen, Annika Rausch, Eva Loth, Bethany Oakley, Tony Charman, Christine Ecker, Declan G. M. Murphy, the EU-AIMS LEAP group, Christian F. Beckmann, Alberto Llera, Jan K. Buitelaar

## Abstract

**Background:** Although many studies have explored atypicalities in gray and white matter (GM, WM) morphology of autism, most of them rely on unimodal analyses that do not benefit from the likelihood that different imaging modalities may reflect common neurobiology. We aimed to establish multimodal brain patterns that differentiate between autism and typically developing (TD) controls and explore associations between these brain patterns and clinical measures.

**Methods:** We studied 183 individuals with autism and 157 TD individuals (6-30 years) in a large deeply phenotyped autism dataset (EU-AIMS LEAP). Linked Independent Component Analysis was utilized to link all participants’ GM and WM images, and group comparisons of modality shared variances were examined. Subsequently, we performed a canonical correlation analysis to explore the aggregated effects between all multimodal GM-WM covariations and clinical profiles.

**Results:** One multimodal pattern was significantly related to autism. This pattern was primarily associated with GM in bilateral insula, frontal, pre- and post-central, cingulate, and caudate areas, and co-occurred with altered WM features in the superior longitudinal fasciculus. The canonical analysis showed a significant multivariate correlation primarily between multimodal brain patterns that involved variation of corpus callosum, and symptoms of social affect in the autism group.

**Conclusions:** Our findings demonstrate the assets of integrated analyses of GM and WM alterations to study the brain mechanisms that underpin autism, and show that the complex clinical autism phenotype can be interpreted by multimodal brain patterns that are spread across the brain involving both cortical and subcortical areas.

## Introduction

Autism Spectrum Disorder (autism) is a heterogeneous condition characterized by difficulties with social and communicative behaviors, repetitive, rigid behaviors and altered sensory processes (1). In search of the brain basis of autism, the condition has been associated with multiple morphological differences in gray matter (GM) and white matter (WM) (2, 3), as reported by magnetic resonance imaging (MRI) studies. However, former studies have shown heterogeneous findings of the alterations in both cortical (e.g., cortical thickness, surface area, volume) and subcortical (e.g., volume) morphometry in multiple brain regions making it difficult to define the neural correlates of autism. For example, researchers found either greater (4) or lower (3) cortical thickness values in temporal areas in autism; and two large-scale studies found different results with respect to the subcortical volume variations (smaller volumes (3) and no difference (5)) in individuals with autism. Additionally, voxel-wise GM volume analyses revealed divergent results in autism, for instance, studies reported increased (6), decreased (7) or unchanged volume (8) in temporal areas. Studies of WM microstructural associations in autism are similarly heterogenous in their findings. Studies have reported that several areas are involved, such as, the corpus collosum (CC), inferior longitudinal fasciculus (ILF) and superior longitudinal fasciculus (SLF) (2, 9, 10). One explanation for discrepant and heterogeneous findings is that the studies differ widely in sample size, sample characteristics, and data analytic strategy - i.e., these studies rely on unimodal analyses techniques that do not benefit from the potentially common neurobiology that different imaging modalities might reflect (11). Additionally, when integrated together these modalities might provide additional analytical sensitivity.

This prompted research to move beyond unimodality and incorporate and connect data from different imaging modalities. For example, (12) suggested that GM variation in autism is generally accompanied by WM variation; (13) showing higher axial diffusivity (L1) in the WM fiber tracts originating and/or terminating in the GM clusters with increased local gyrification in adults with autism. Despite the progress away from unimodal approaches, in essence, these MRI studies which correlate GM and WM measures do so after separate unimodal statistical analyses. This likely has less sensitivity to assess the biological variance than fully integrating multimodal data analysis across participants.

Here, we aim to utilize an integrative multivariate approach, linked independent component analysis (LICA), to simultaneously incorporate several imaging modalities allowing the investigation of inter-subject variability across modalities in one analysis (14, 15). So far, studies that highlight the underlying shared variance between modalities using LICA in autism remain scarce. Previous studies revealed case-control differences between adults with autism and typically developing (TD) individuals in linked voxel-based morphometry (VBM) and diffusion tensor imaging (DTI) measures patterns in several brain regions (16, 17). However, these studies focused exclusively on adult individuals with high functioning autism and were comprised of relatively small sample sizes (<100 individuals) (16, 17). Additionally, the study by (16) only included male participants. Autism is a highly diverse condition; we therefore investigate multimodal patterns in a broader more representative autism sample which might help better characterize brain patterns of autism - one of the aims of the current study.

In addition to identifying categorical group differences, dimensional analyses, i.e., analyses of continuous scores of autism symptoms might capture more of the heterogeneity of autism compared to categorical diagnostic labels. Many studies have demonstrated the univariate connections between GM or WM patterns and the core symptoms of autism (e.g., (2, 3)). Nonetheless, the relationships between brain substrates and clinical phenotypes are potentially the outcome of sintegrative effects across multiple autism symptom domains and brain areas, and therefore the multidimensional associations between brain GM-WM covariations and core symptoms of autism need to be clarified. Consequently, canonical correlation analysis (CCA), as a multivariate approach, is effective to learn such associations from a more comprehensive perspective in autism (18).

This study was designed to overcome the aforementioned limitations of previous work by applying LICA to the Longitudinal European Autism Project (LEAP) dataset (19) to link GM and WM sources of variance. The LEAP dataset provides a deeply phenotyped and comprehensively biologically assessed multisite sample of individuals with/without autism that allows relating the results of LICA to clinical characteristics of the participants. More specifically, we applied (a) a univariate approach to identify categorical group difference of linking GM-WM brain patterns, and subsequently their one-to-one relations to continuous measures of autism symptoms; (b) a multivariate method (i.e., CCA) to further quantify the association between two datasets of brain inter-modalities patterns and autism symptoms in the autism group.

## Methods and Materials

### Participants

The participants were part of the EU-AIMS and AIMS-2-TRIALS Longitudinal European Autism Project (LEAP) dataset - a large multicenter study aimed at identifying and validating biomarkers in autism (19, 20). Individuals with autism were included based on an existing clinical diagnosis according to DSM-IV, DSM-IV-TR, DSM-5, or ICD-10. Each participant underwent clinical, cognitive, and MRI assessment at one of six collaborative centers. We refer to (19, 20) for further details on experimental design and clinical characterization. In the present study, diffusion-weighted image (DWI) data at timepoint 1 were only available from participants in three centers. Therefore, the participants were selected who had both T1-weighted and DWI data available from the following centers: Institute of Psychiatry, Psychology and Neuroscience, King’s College London, United Kingdom; Radboud University Medical Centre, Nijmegen, the Netherlands; Central Institute of Mental Health, Mannheim, Germany (Supplementary Table S1).

### Clinical measures

The Autism Diagnostic Interview-Revised (ADI) (21) and the Autism Diagnostic Observational Schedule 2 (ADOS) (22) were used to measure the past (ever and previous 4-to-5 years) and current core symptom severities of autism from social interaction, communication, and restricted repetitive behaviors (RRB) domains. Additionally, we used several parent-reported scales to assess autism symptoms, including the Social Responsiveness Scale 2nd Edition (SRS) (23) capturing the social-communication variations, the Repetitive Behavior Scale-Revised (RBS) (24) identifying the repetitive and rigid behaviors, and the Short Sensory Profile (SSP) (25) evaluating the sensory processing variations. Concerning the potential effect of Attention Deficit Hyperactivity Disorder (ADHD) co-occurrence, we included the two dimensions (inattention and hyperactivity/impulsivity symptoms) of ADHD DSM-5 rating scale (26) as the additional covariates in the post-hoc analyses. The ADHD rating scale we used was based on parent-report. There was a substantial amount of missing clinical data (e.g., for the SSP only 108 out of 185 participants had available data in the autism group) which could greatly reduce the power of our analysis. To tackle the missing clinical data and fully harness the large LEAP sample size we used imputed clinical data (27).

### MRI data acquisition

All participants were scanned on 3T MRI scanners. High-resolution structural T1-weighted images were acquired using magnetization-prepared rapid gradient-echo sequence with full head coverage, at 1.2 mm thickness with 1.1×1.1 mm in-plane resolution. Diffusion-weighted imaging (DWI) scans were acquired using echo-planar imaging sequence, at 2 mm thickness with 2.0×2.0 mm in-plane resolution.

MRI data acquisition parameters can be found in the Supplementary Table S2.

## Image processing

### GM volume estimation

Structural T1 images were preprocessed according to CAT12 toolbox (https://dbm.neuro.uni-jena.de/cat/) pipeline in SPM12 (Wellcome Department of Imaging Neuroscience, London, UK) to obtain VBM data, which is a spatially-unbiased whole-brain approach extracting voxel-wise GM volume estimations. T1-weighted images were automatically segmented into GM, WM, and cerebrospinal fluid and affine registered to the MNI template. A high-dimensional, nonlinear diffeomorphic registration algorithm (DARTEL) (28) was used to generate a study-specific template from GM and WM tissue segments of all participants, and then to normalize all segmented GM maps to MNI space with 2mm isotropic resolution. All GM images were smoothed with a 4mm full-width half-max (FWHM) isotropic Gaussian kernel.

### Diffusion parameters

DWI images from all sites were preprocessed using the same pipeline. De-nosing was performed using the Marchenko-Pastur principal component analysis (MP-PCA) method (29). Subsequently, Gibbs-ringing artefacts were removed according to (30). FSL *eddy* was employed to correct the eddy-current induced distortions and subject motion (31). To improve the final quality of data and recover most the motion artefacts, we utilized intra-volume slice motion correction (32). Quality control reports were then generated for each subject and each site (33).

Individual voxel-wise FA, mean diffusivity (MD), mode of anisotropy (MO), L1 and radial diffusivity (RA) maps were derived using dtifit in FSL (34). FA images were processed using Tract-Based Spatial Statistics (TBSS) pipeline including registration of all images to FMRIB58_FA standard space, skeletonization of the mean group white matter and projection of each individual’s data onto the skeleton (35). The mean skeleton image was thresholded at FA 0.2. Other DTI measures (MD, MO, L1, RD) were projected onto the FA skeleton using the tbss_non_FA option. All DTI data had 1mm isotropic resolution when entering the following data fusion model.

A full quality control report and additional preprocessing details of the GM and WM images are included in the Supplementary Section 3.

### Modalities fusing analysis

The shared inter-participant variations across six features (i.e., VBM, FA, MD, MO, L1, RD) were explored using LICA (11). LICA is able to factorize the multiple input modalities simultaneously into modality-wise independent components (ICs) while importantly constraining all decompositions to be linked through a shared participant-loading matrix, which describes the amount of contribution of each participant to a specific IC. In addition to the participant-loading matrix, this method provides, per IC, a vector reflecting the contribution (weight) of each modality and a spatial map per modality showing the extent of the spatial variation. All mathematical algorithms of LICA are detailed in (11). As the model order is recommended to be less than 25% of the sample size (11, 14), 80-dimensional factorization was chosen to perform LICA. A multimodal index (36) was calculated to present the contribution uniformity of the modalities in each IC, in which a value of 1 denotes the involved modalities contributing equally to the given IC, and a value close to 0 means one modality dominating the IC variability. Note, our study aimed at detecting autism-related multimodal inter-subject variations, we therefore excluded the ICs comprising one single modality weighted more than 50% (36).

### Statistical approach

The participant-loadings characterize the inter-individual variations of the multi-modal effects, and in the current study, they were used for the analyses of group differences between autistic and TD individuals, and for associations with behavioral measures. Results reported in the main text are performed using imputed data to maximize the statistical power. All analyses were replicated using the original non-imputed data (Supplementary Section 4).

### Case-control difference

A generalized linear model (GLM) was utilized to examine group differences of the brain’s inter-participant variations in multimodal (i.e., no single modality contributed more than 50%) LICA outputs while controlling for age, sex, IQ and scanner site. Multiple comparison correction was implemented using false discovery rate (FDR) (p<0.05) (37).

### Brain-behavior associations

Similarly, we used a GLM to explore the univariate associations between each multimodal IC and subscales of ADI and ADOS, SRS, RBS, and SSP in the autism group while controlling for age, sex, IQ and scanner site. We corrected for multiple comparisons with FDR (p<0.05). Subsequently, we utilized CCA (18) to better picture the overall association between all multimodal brain ICs and all symptom phenotypes in the autism group. In this study, we referred to each pair of canonical variates as CCA mode. The statistical significance of CCA modes was assessed by permutation inference (38). Since the behavior profiles were evaluated by either qualified examiners or parent-reporting, we performed two separate CCA analyses on the basis of the assessment type to relate multimodal ICs to subsets of behavioral measures; in the first CCA analyses (CCA_1_) we included the subscales of ADI and ADOS, which were rated by qualified examiners, while in the second (CCA_2_) we used total scores of parent-rated SRS, RBS, and SSP within autistic individuals. For each CCA’s multiple testing correction, we used stepwise cumulative maximum approach, p<0.05, see details in (38). The evaluation of the contribution of each IC and each clinical measure to the canonical correlation was according to the structural coefficient of each variable described previously (39).

## Results

The quality of all raw T1, DWI and preprocessed data was carefully checked resulting in the exclusion of 4 individuals based on the presence of structural abnormalities, visible artifacts, or preprocessing failures (for details, see Supplementary Section 3). This resulted in a final sample of 344 participants, including 185 individuals with autism and 159 TD individuals. The demographic and clinical information of the final sample is summarized in Table 1.

**Table 1.**
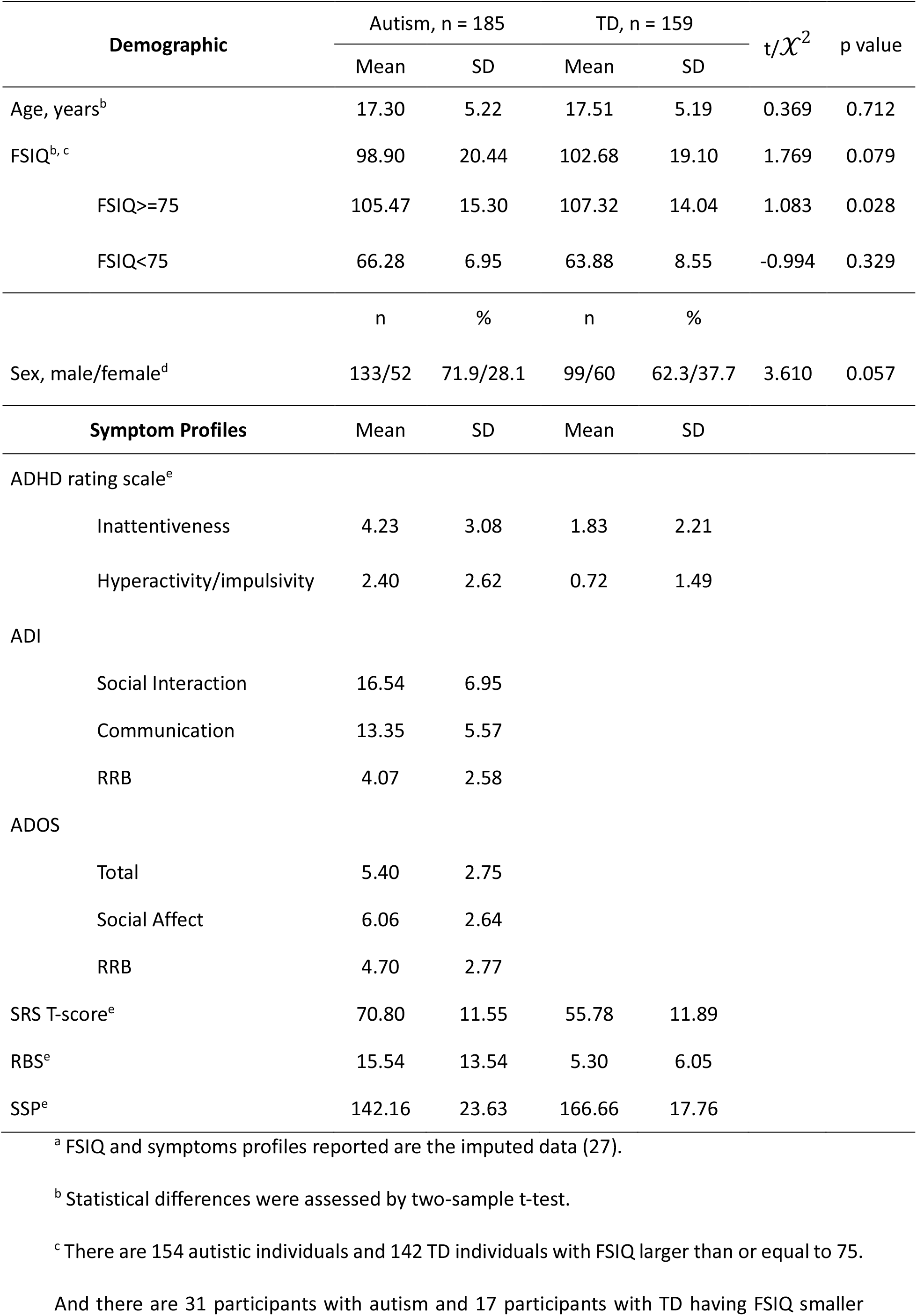
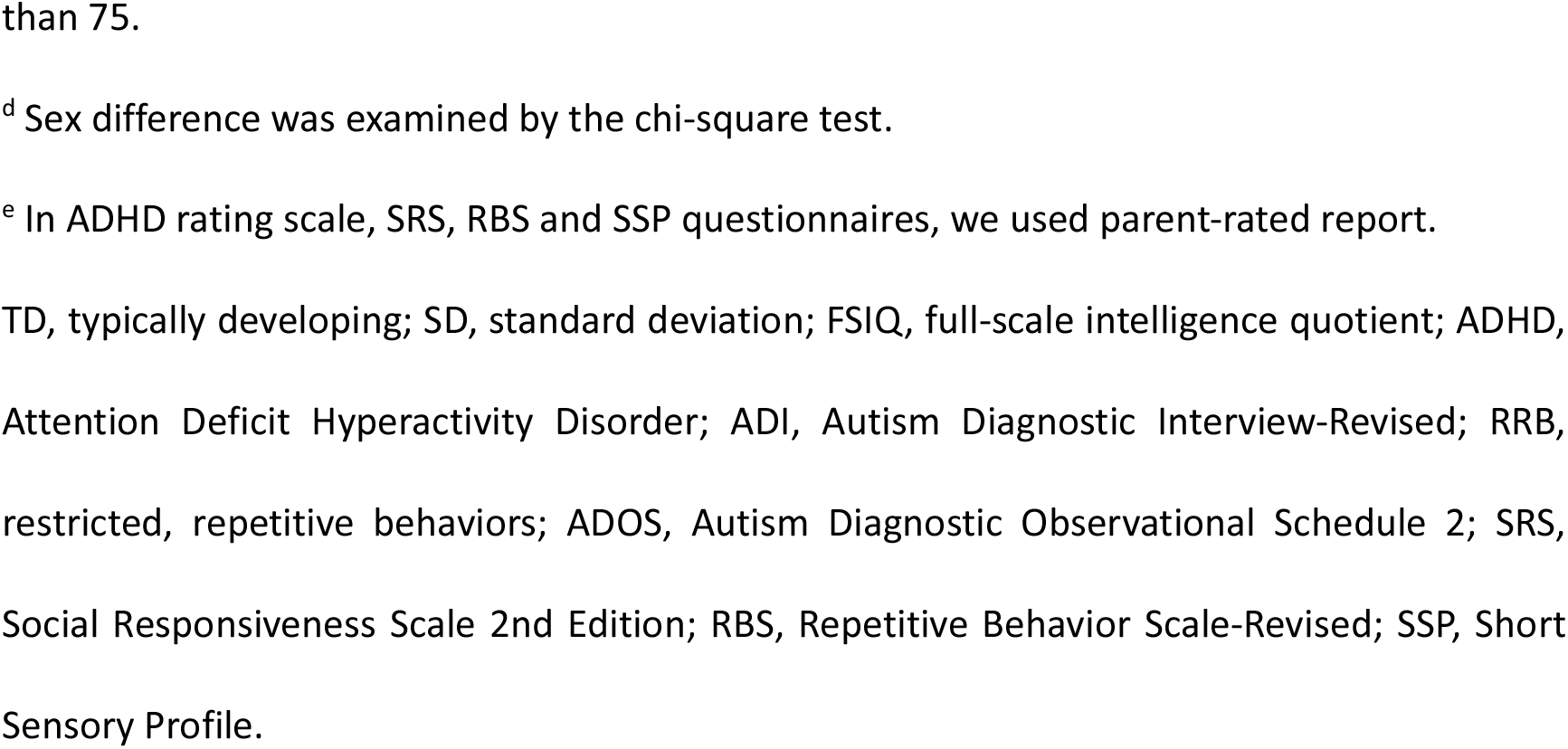
Demographic information of participants^a^

| Demographic | Autism, n = 185 | | TD, n = 159 | | t/$\chi^2$ | p value |
| --- | --- | --- | --- | --- | --- | --- |
|  | Mean | SD | Mean | SD |  |  |
| Age, years<sup>b</sup> | 17.30 | 5.22 | 17.51 | 5.19 | 0.369 | 0.712 |
| FSIQ<sup>b, c</sup> | 98.90 | 20.44 | 102.68 | 19.10 | 1.769 | 0.079 |
| FSIQ≥75 | 105.47 | 15.30 | 107.32 | 14.04 | 1.083 | 0.028 |
| FSIQ<75 | 66.28 | 6.95 | 63.88 | 8.55 | -0.994 | 0.329 |
|  | n | % | n | % |  |  |
| Sex, male/female<sup>d</sup> | 133/52 | 71.9/28.1 | 99/60 | 62.3/37.7 | 3.610 | 0.057 |
| **Symptom Profiles** | Mean | SD | Mean | SD |  |  |
| ADHD rating scale<sup>e</sup> |  |  |  |  |  |  |
| Inattentiveness | 4.23 | 3.08 | 1.83 | 2.21 |  |  |
| Hyperactivity/impulsivity | 2.40 | 2.62 | 0.72 | 1.49 |  |  |
| ADI |  |  |  |  |  |  |
| Social Interaction | 16.54 | 6.95 |  |  |  |  |
| Communication | 13.35 | 5.57 |  |  |  |  |
| RRB | 4.07 | 2.58 |  |  |  |  |
| ADOS |  |  |  |  |  |  |
| Total | 5.40 | 2.75 |  |  |  |  |
| Social Affect | 6.06 | 2.64 |  |  |  |  |
| RRB | 4.70 | 2.77 |  |  |  |  |
| SRS T-score<sup>e</sup> | 70.80 | 11.55 | 55.78 | 11.89 |  |  |
| RBS<sup>e</sup> | 15.54 | 13.54 | 5.30 | 6.05 |  |  |
| SSP<sup>e</sup> | 142.16 | 23.63 | 166.66 | 17.76 |  |  |

<sup>a</sup> FSIQ and symptoms profiles reported are the imputed data (27).
<sup>b</sup> Statistical differences were assessed by two-sample t-test.
<sup>c</sup> There are 154 autistic individuals and 142 TD individuals with FSIQ larger than or equal to 75.
And there are 31 participants with autism and 17 participants with TD having FSIQ smaller
than 75.
<sup>d</sup> Sex difference was examined by the chi-square test.
<sup>e</sup> In ADHD rating scale, SRS, RBS and SSP questionnaires, we used parent-rated report.
TD, typically developing; SD, standard deviation; FSIQ, full-scale intelligence quotient; ADHD, Attention Deficit Hyperactivity Disorder; ADI, Autism Diagnostic Interview-Revised; RRB, restricted, repetitive behaviors; ADOS, Autism Diagnostic Observational Schedule 2; SRS, Social Responsiveness Scale 2nd Edition; RBS, Repetitive Behavior Scale-Revised; SSP, Short Sensory Profile.

### Group effect of multimodal integration components

We obtained 80 ICs from the multimodal integration analysis, 75 ICs of which were identified as multimodal components (i.e., no single modality weighted more than 50%). The modality contributions (for 80 ICs) and multimodal index of each IC can be found in Supplementary Figure S3. We subsequently used the participant-loadings of the 75 ICs to test for group differences and found one component (IC58) with a significant case-control difference (β=-0.192, FDR corrected p=0.028; Figure 1). The respective contributions of the modalities in IC58 are 26% from VBM, 18% from FA, 18% from MO, 14% from L1, 14% from RD, and 10% from MD, indicating that various MRI features share variance associated with autism. In Figure 1, we present the summarized images of each modality’s spatial map of IC58. The spatial patterns show autism-related smaller GM volume in the bilateral insula, inferior frontal gyrus (IFG), orbitofrontal cortex (OFC), precentral, postcentral gyrus, lateral occipital cortex (LOC), inferior temporal gyrus (ITG), angular gyrus (AG), posterior division of cingulate gyrus (PCC), and precuneus cortex, and larger GM volume in calcarine cortex, bilateral middle frontal gyrus (MFG), caudate and anterior division of cingulate gyrus (ACC). Correspondingly, autism-related DTI features were found in bilateral SLF, corticospinal tract (CST), and inferior fronto-occipital fasciculus (IFOF). In addition to these fasciculi, RD and MD in the cingulum and anterior thalamic radiation were also implicated. Taken together, the implication of superior longitudinal fasciculi and their adjacent GM volumes; frontal, precentral, and postcentral areas (Supplementary Figure S4) in autism indicate that variations of GM volumes and WM microstructure are linked in these brain locations, rather than modality or tissue dependent. These results were not significantly driven by scan site (Supplementary Section 7).

**Figure 1.**
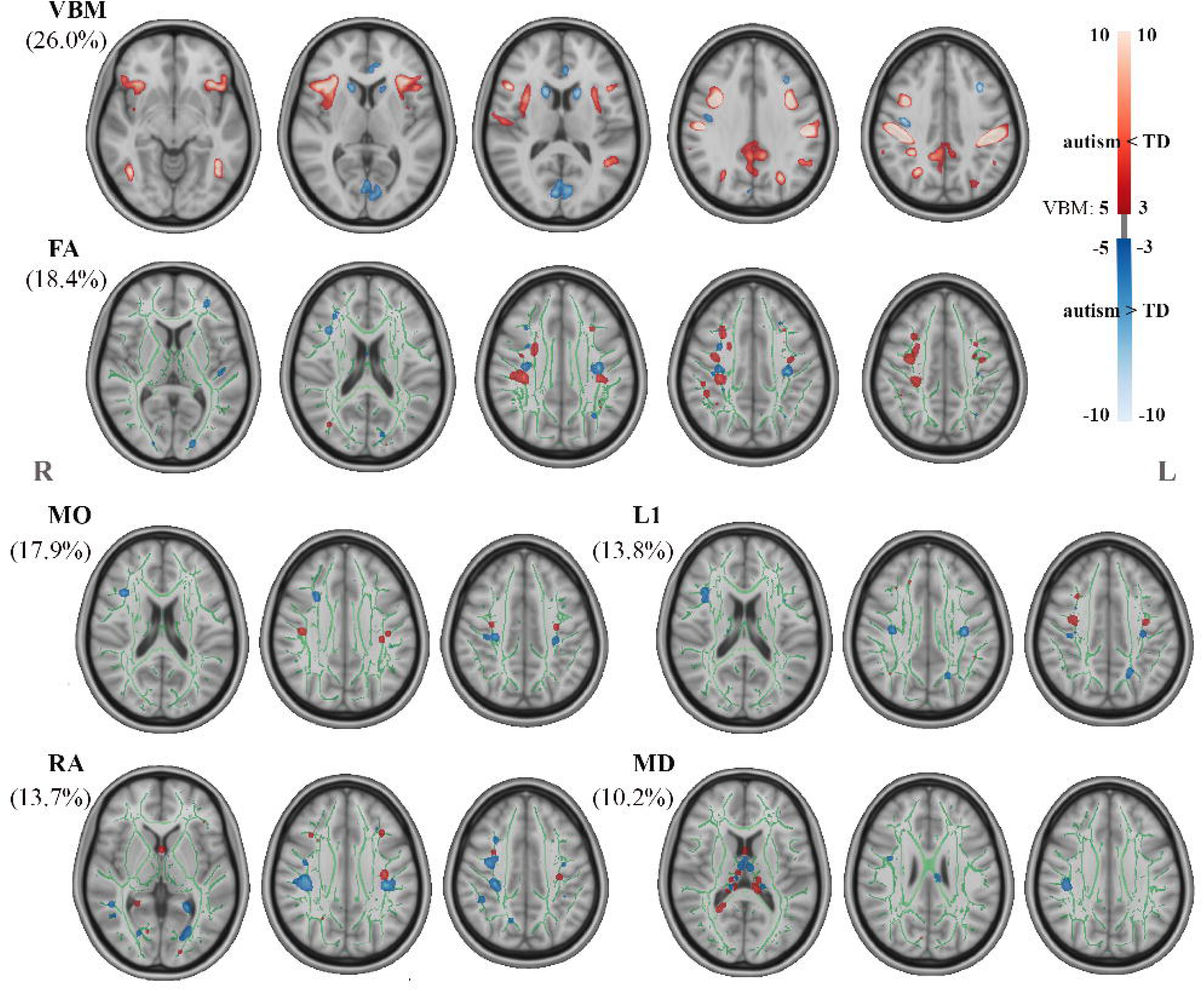
The multimodal component shows significant case-control difference. The relative contribution of each feature is displayed in brackets. The VBM spatial map is thresholded at 5<|Z|<10. Clusters of DTI features were filled and thresholded at 3<|Z|<10, then smoothed using a 0.3mm Gaussian kernel in FSL for visualization purposes. VBM, voxel-based morphometry; FA, fractional anisotropy; MO, mode of anisotropy; L1, axial diffusivity; RA, radial diffusivity; MD, mean diffusivity; TD, typically developing.

Post-hoc, to assess the influence of co-occurring ADHD symptoms on the multimodal IC found significantly associated with group, we additionally included inattention and hyperactivity/impulsivity scores of parent-reported ADHD rating scale (26) as covariates in the GLM of IC58. The analysis showed that the group effect of IC58 was also robust to the inclusion of continuous scores of inattention and hyperactivity/impulsivity symptoms as additional covariates in the model (β=-0.192, p=0.002).

### Relating multimodal integration pattern to behavior profiles

We conducted the univariate (GLM) and multivariate (CCA) correlation analyses on brain and behavior data in the autism group only. No significant univariate brain-behavior relationship in the autism group were found (FDR corrected p>0.300). We did however find a significant multivariate association pattern of CCA_1_ (linking ADI and ADOS subscales to multimodal ICs) (r=0.788, corrected p=0.002, Figure 2). In this multivariate associated pattern, multimodal IC1 (canonical weight: 0.407) and IC33 (canonical weight: −0.096) showed the strong contribution to the correlation with autism core symptoms, while from a phenotypic perspective this multivariate pattern demonstrated a strong association with the ADOS social affect (SA) and RRB subscales. WM microstructure mainly dominated in IC1 and IC33. IC1 primarily involved shared variations in CC, bilateral anterior thalamic radiation and CST, while IC33 was governed by variation in CC. These two predominant ICs highlight the involvement of CC in autism symptoms.

**Figure 2.**
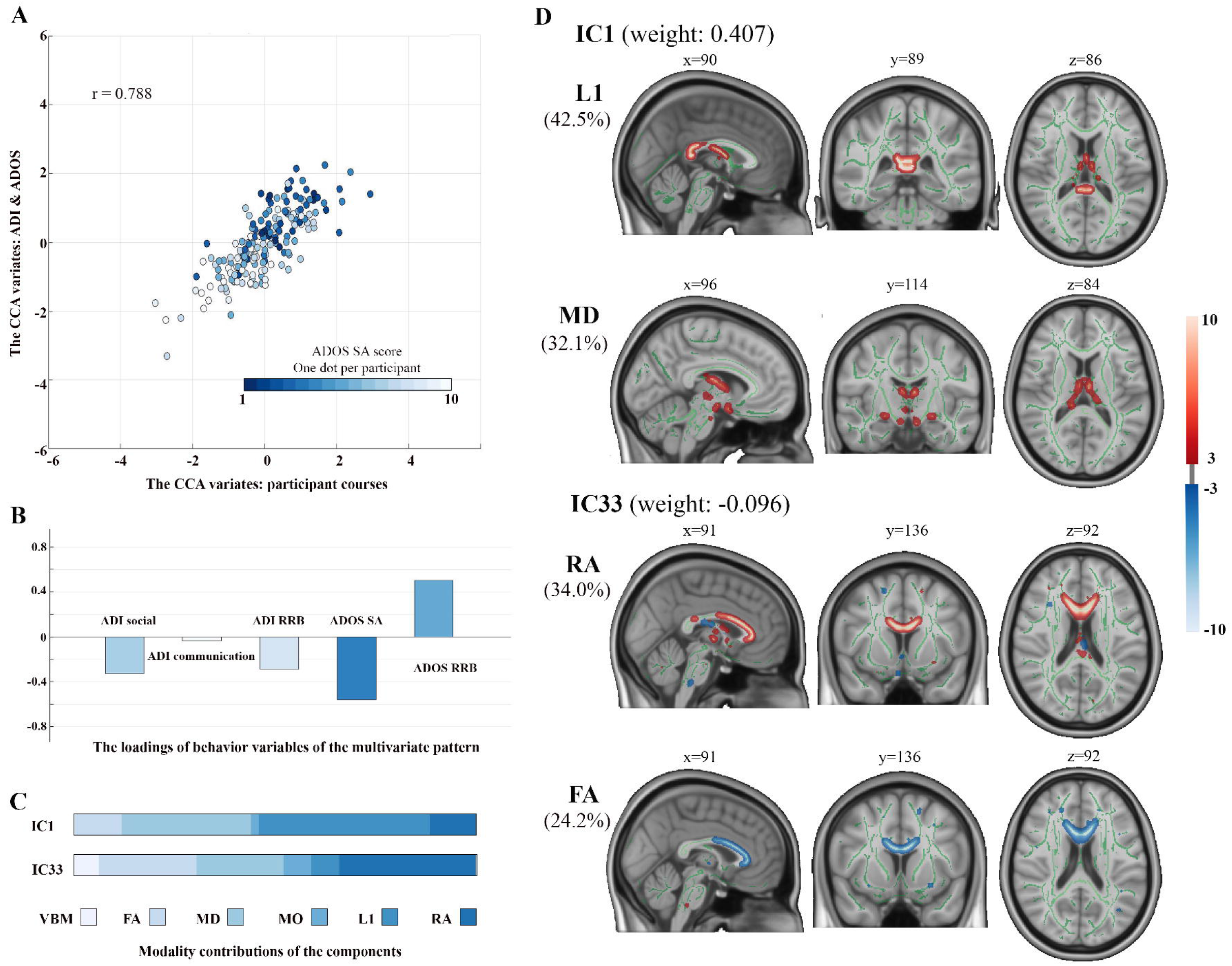
The multivariate association pattern is found significant between multimodal components and subscales of ADI and ADOS. **A** displays the scatterplot of this association pattern, and x, y axes are the pair of variates that derives from CCA. One dot in each participant is coded with gradient color regarding to the SA subscale of ADOS. **B** demonstrates the loading of each behavior subscale in this association pattern. **C** shows the modality contributions of the multimodal components displayed in **D. D** exhibits the two multimodal components contributed most to the association pattern. The weights of each component are showed in the brackets. The modality spatial maps are thresholded at 3<|Z|<10. All CCAs were only performed in autism group. CCA, canonical correlation analysis; ADI, Autism Diagnostic Interview-Revised; ADOS, Autism Diagnostic Observational Schedule 2; SA, social affect; RRB, restricted repetitive behavior; IC, independent component; MO, mode of anisotropy; RA, radial diffusivity; MD, mean diffusivity; FA, fractional anisotropy; L1, axial diffusivity; VBM, voxel-based morphometry.

## Discussion

We examined autism-related inter-individual variance of integrated GM and WM morphology in a large representative sample of individuals with and without autism. Analyses showed a significant diagnostic-group effect of the linked GM-WM pattern that supports our hypothesis of the link between GM and WM morphology alterations in individuals with autism. In particular, the GM volume variation in pre- and post-central areas converged with the WM microstructural variation in the SLF. This spotlights the shared variances between GM and WM morphology in these brain areas in autism, and suggests the structural associations in autism are not only limited to localized regions but also involve the WM pathways connecting these brain areas. In a next set of analyses, we found a significant integrative association between brain multimodal patterns and autism core symptoms using CCA in the autism group, where the identified brain multimodal patterns underline the important role of WM morphology, particularly the involvement of CC.

Notably, the autism-specific VBM pattern on this multimodal analysis corroborates our previous unimodal GM volume covariation study in a larger sample of the EU-AIMS project (8). The areas of bilateral insula, IFG, OFC, and caudate form a steady autism-related covariation pattern in previous and current studies. These areas were demonstrated previously to relate to repetitive behaviors and reward-based decision-making abilities in autism (40, 41). The covariation of insula and frontal areas in our studies indicates the consistency and stability of the co-occurring GM morphological alterations in autism. Moreover, in agreement with the effectiveness of multimodal/multivariate approaches suggested by previous studies (42, 43), the application of the LICA approach provides increased sensitivity to detect autism-related brain regions. In our study, this extends identified GM associations to pre-central, post-central, occipital and temporal areas compared to unimodal analysis, and incorporates significant WM findings of DTI measures. These otherwise unrecognized small effects in each modality were detected by modeling the variances across modalities. In deviance from our previous study (Mei et al., 2020), the current results did not include significant autism-related alterations of limbic-crucial areas (amygdala and hippocampus), which might be attributed to the smaller sample size.

Our results indicated one covarying set of brain GM and WM areas associated with autism diagnosis. In this multimodal set, GM volume in cortical and subcortical regions and microstructure in WM tracts (mainly SLF, CST and IFOF) were implicated and these regions/tracts have previously been identified in unimodal analyses (10, 40, 44–46). This broad range of brain regions along with large WM bundles associated with autism is in accord with the notion that the neural correlates of autism are widespread in brain regions and connectivity patterns (47–49). This also corresponds with another multimodal autism study reporting extensive autism-related brain areas (16). The areas of this IC have been linked previously to both social and non-social cognitive difficulties in individuals with autism, varying from visual, sensory and motor processing to high-order cognitive abilities (10, 50–53). For example, pre-central, post-central gyrus, SLF and CST are related to (sensory-)motor processing and have been implicated in autism (10, 46, 54). Additionally, the LOC and IFOF are two areas implicated in varied visuospatial processing in autism (51, 55). These adjacent affected areas (grouped areas of pre-, post-central areas and SLF, CST; grouped areas of LOC and IFOF occipital section) in our findings logically is in line with the brain organization principles during development, which states that nearby areas tend to be more interconnected (56, 57). In summary, the autism diagnosis-related co-varying GM-WM pattern reflect that autism is a complex condition associated with neural morphology. However, we did not find any significant univariate relationship between behavioral phenotypes and GM-WM patterns. This is probably a result of the diverse phenotypes in our sample (i.e., complex and heterogenous nature of autism), therefore, the compound variances of the symptom profiles cannot be explained by single multimodal brain patterns. Additionally, imaging studies suggested that individuals with autism develop alternative processing strategies (48) that might mix or neutralize the manifestations of behavioral phenotypes in autism moderating detection of well-established brain-behavior relations.

Although we do not observe an association between autism diagnosis-related IC and symptom/behavior profiles, there is one prominent WM dominated multivariate relation between all multimodal brain patterns and subscales of ADI and ADOS. The top two ranking ICs emphasize the importance of WM connection to the core traits of autism, especially the microstructure of CC. Multivariate/multimodal analysis increase the difficulty in interpreting findings, as it’s challenging to clarify the direction of each association. Nonetheless, coinciding with previous studies (2, 58), the alterations of CC microstructural measures are evidently associated with more severe core symptoms in autism, which were frequently reported in previous autism research, including its genu, body and splenium parts (10, 59, 60). Significantly, GM volume contributed only by a small amount to the multivariate correlation, which implicates WM morphology has a stronger connection to the autism behavioral phenotypes compared to GM. In our previous GM work a multivariate correlation pattern exhibited a strong association between RRB scores of ADI and ADOS and GM covariations in autism, while here when including WM microstructural measures, the multimodal brain patterns demonstrated a strong association with SA and RRB domains of the ADOS. This multivariate brain-behavior association needs further investigation to determine the relationship between the development of WM microstructure and behaviors, which might expand our knowledge of current brain-behavior association patterns.

Our findings should be interpreted with regard to several limitations. First, to generalize our pattern of brain alterations associated with autism requires replication in other large-scale datasets. Second, the current multimodal data set included fewer participants than our previous work (8), which may have lowered statistical power when detecting the group effects and brain-behavior associations in autism group. Despite that, this is still the largest multimodal MRI study of autism to date and includes a diverse sample of autistic and neurotypical participants.

In current study, we demonstrate autism-related inter-individual covariations of GM volume in frontal, pre-central, post-central and occipital areas and microstructure in associated WM fasciculi. Together, these GM and WM alterations are part of the underlying neural substrates of the phenotypes in autism. Subsequently, we highlight the potential role of WM, specifically CC, in the relation to the core symptoms of autism. Further studies may link our GM-WM morphometric findings with brain function acquired from cognitive assessments and/or functional MRI data.

## Acknowledgments

This work is primarily supported by the EU-AIMS consortium (European Autism Interventions), which receives support from Innovative Medicines Initiative Joint Undertaking Grant No.115300, the resources of which are composed of financial contributions from the European Union’s Seventh Framework Programme (Grant No. FP7/2007-2013), from the European Federation of Pharmaceutical Industries and Associations companies’ in-kind contributions; and by the AIMS-2-TRIALS consortium (Autism Innovative Medicine Studies-2-Trials), which has received funding from the Innovative Medicines Initiative 2 Joint Undertaking under grant agreement No. 777394, and this Joint Undertaking receives support from the European Union's Horizon 2020 research and innovation programme and EFPIA and AUTISM SPEAKS, Autistica, SFARI. This work reflects the authors’ views and neither IMI nor the European Union, EFPIA or any Associated Partners are responsible for any use that may be made of the information contained therein.

TM is supported by a China Scholarship Council grant (No 201806010408). DLF is supported by funding from the European Union’s Horizon 2020 research and innovation programme under the Marie Skłodowska-Curie grant agreement No 101025785. This work has been further supported by the European Union Seventh Framework Programme Grant Nos. 602805 (AGGRESSOTYPE) (to JKB), 603016 (MATRICS) (to JKB), and 278948 (TACTICS) (to JKB); European Community’s Horizon 2020 Programme (H2020/2014-2020) Grant Nos. 643051 (MiND) (to JKB), 642996 (BRAINVIEW) (to JKB) and 847818 (CANDY) (to JKB and CFB); the Netherlands Organization for Scientific Research VICI Grant No. 2020/TTW/00836465 (to CFB); Wellcome Trust Collaborative Award Grant No. 215573/Z/19/Z (to CFB); the Autism Research Trust (to SBC).

We thank all participants and their families for participating in this study. We gratefully acknowledge the contributions of all members of the EU-AIMS LEAP group: Jan K. Buitelaar (primary contact/principal investigator), Jumana Ahmad, Sara Ambrosino, Bonnie Auyeung, Tobias Banaschewski, Simon Baron-Cohen, Sarah Baumeister, Christian F. Beckmann, Sven Bölte, Thomas Bourgeron, Carsten Bours, Michael Brammer, Daniel Brandeis, Claudia Brogna, Yvette de Bruijn, Bhismadev Chakrabarti, Tony Charman, Ineke Cornelissen, Daisy Crawley, Flavio Dell’Acqua, Guillaume Dumas, Sarah Durston, Christine Ecker, Jessica Faulkner, Vincent Frouin, Pilar Garcés, David Goyard, Lindsay Ham, Hannah Hayward, Joerg Hipp, Rosemary Holt, Mark H. Johnson, Emily J.H. Jones, Prantik Kundu, Meng-Chuan Lai, Xavier Liogier D’ardhuy, Michael V. Lombardo, Eva Loth, David J. Lythgoe, René Mandl, Andre Marquand, Luke Mason, Maarten Mennes, Andreas Meyer-Lindenberg, Carolin Moessnang, Nico Mueller, Declan G.M. Murphy, Bethany Oakley, Laurence O’Dwyer, Marianne Oldehinkel, Bob Oranje, Gahan Pandina, Antonio M. Persico, Annika Rausch, Barbara Ruggeri, Amber Ruigrok, Jessica Sabet, Roberto Sacco, Antonia San José Cáceres, Emily Simonoff, Will Spooren, Julian Tillmann, Roberto Toro, Heike Tost, Jack Waldman, Steve C.R. Williams, Caroline Wooldridge, Iva Ilioska, Ting Mei and Marcel P. Zwiers.

## Disclosures

TC has received consultancy from Roche and Servier and received book royalties from Guildford Press and Sage. DGM has been a consultant to, and advisory board member, for Roche and Servier. He is not an employee of any of these companies, and not a stock shareholder of any of these companies. CFB is director and shareholder in SBGNeuro Ltd. TB served in an advisory or consultancy role for ADHS digital, Infectopharm, Lundbeck, Medice, Neurim Pharmaceuticals, Oberberg GmbH, Roche, and Takeda. He received conference support or speaker’s fee by Medice and Takeda. He received royalities from Hogrefe, Kohlhammer, CIP Medien, and Oxford University Press. JKB has been a consultant to, advisory board member of, and a speaker for Janssen Cilag BV, Eli Lilly, Shire, Lundbeck, Roche, and Servier. He is not an employee of any of these companies, and not a stock shareholder of any of these companies. He has no other financial or material support, including expert testimony, patents or royalties. The present work is unrelated to the above grants and relationships. The other authors report no biomedical financial interests or potential conflicts of interest.

