## Supplementary Information for "Autism is associated with inter-individual variations of gray and white matter morphology"

### 1. Demographic information of each site

**Table S1**. Demographic information of participants in each site

| Variable | KCL | | Nijmegen | | Mannheim | |
|---|---|---|---|---|---|---|
| | Autism | TD | Autism | TD | Autism | TD |
| N | 91 | 71 | 72 | 55 | 22 | 33 |
| Age,mean,[SD] | 17.87 [5.24] | 16.69 [5.84] | 16.95 [5.62] | 15.92 [4.08] | 16.07 [2.86] | 15.46 [3.03] |
| FIQ,mean,[SD] | 99.04[22.19] | 104.48[22.70] | 97.87[19.15] | 97.14[15.29] | 101.73[15.71] | 108.02[12.71] |
| Sex, N, [%] | | | | | | |
| Male | 64 [70.33] | 40 [56.34] | 52 [72.22] | 36 [65.45] | 17 [77.27] | 23 [69.70] |

TD, typically developed; SD, standard deviation; FIQ, full-scale intelligence quotient.

### 2. MRI data acquisition parameters

**Table S2**. Summary of acquisition parameters across sites

| | KCL | Nijmegen | Mannheim |
|---|---|---|---|
| **MPRAGE T1 weighted sequence** | | | |
| Manufacturer | GE Medical systems | Siemens | Siemens |
| Model | Discovery mr750 | Skyra | TimTrio |
| Software Version | LX MR DV23.1_V02_1317.c | Syngo MRD13 | Syngo MR B17 |
| Acquisition sequence | SAG ADNI GO ACC SPG | Tfl3d1_16ns | MPRAGE ADNI |
| Coverage | 256*256 | 256*256 | 256*256 |
| slices | 196 | 176 | 176 |
| Thickness [mm] | 1.2 | 1.2 | 1.2 |
| Resolution [$mm^3$] | 1.1*1.1*1.2 | 1.1*1.1*1.2 | 1.1*1.1*1.2 |
| TR [s] | 7.31 | 2.3 | 2.3 |
| TE [ms] | 3.02 | 2.93 | 2.93 |
| FA [°] | 11 | 9 | 9 |
| FOV | 270 | 270 | 270 |
| **EPI diffusion weighted sequence** | | | |
| TR / TE (ms) | 1200 / 67 | 1200 / 70 | 1200 / 96 |
| Flip angle (°) | 90 | 90 | 90 |
| Slice thickness (mm) / No. | 2 / 72 | 2 / 72 | 2 / 72 |
| In-plane resolution (mm2) | 2 x 2 | 2 x 2 | 2 x 2 |
| B-values (s/mm2) | 0 / 1500 | 0 / 1500 | 0 / 1500 |
| No. of gradients | 6 / 60 | 6 / 60 | 6 / 60 |

### 3. Quality control report

The current work proceeded our previous work (1), where we used 604 participants with preprocessed voxel-based morphometry (VBM) data. In the previous work we excluded those without FIQ (n=5) however here we use imputed demographic and clinical data and therefore include those participants. New participants were additionally added (N=96). This gave us 700 subjects in total with VBM data. However, in our LEAP wave 1 dataset, only 3 scanning sites had diffusion weighted imaging (DWI) data of appropriate quality limiting our total sample to 418 participants.

Quality control reports of preprocessing of DWI data were generated for each participant and each site (three sites included: KCL, Nijmegen and Mannheim) (2). 49 participants with excessive motion during acquisition (absolute motion > 4 mm translation), with high number of outliers (>6%) were excluded from the study. To also correct for possible signal drop-out in the b0 data, b0 voxels for each slice were compared and outliers were identified using the conventional interquartile range (IQR) method for outlier detection (outlier threshold=3$^{rd}$ quartile+1.5xIQR). Slices that were considered outliers were then replaced with the corresponding median b0. Finally, to make the diffusion data more uniform across sites, all data was resampled using a real symmetric spherical harmonics (SH) representation of the DWI signal limited to SH degrees l=6. This allowed a further reduction of high frequency noise and allow the detection and replacement of remaining data outliers not fully recovered by eddy. Because some of the datasets from Siemen's scanners still exhibited residual Gibbs ringing in b0s data compared to the other sites, a final Gibbs ringing correction was applied to all datasets using ExploreDTI only to b0 data (3). After visual inspection data quality looked consistent across datasets and with no visible residual artefacts (there were 18 participants excluded due to artifacts or bad coverage of brain). Please note, because Nijmegen and Mannheim exhibited a high Rician noise floor in the raw data (4), before running the main pre-processing pipeline, datasets from these sites were pre-corrected using an in-house implementation of the methods described in (5) to reduce the bias in the DWI signal induced by the noise floor. The above procedures resulted in 351 participants in total with high quality DWI data, in which 3 participants without T1 images were excluded, therefore there were 348 participants entering VBM and DTI generations.

A VBM quality control report was generated (based on the participants with both T1 and DWI images) by the CAT SPM pipeline for each participant that included visual evaluation of the segmentation, and quantitative quality measures including mean correlation from sample homogeneity module and weighted overall image quality ratings that were additionally used to detect and exclude images of insufficient quality for inclusion in analysis. Accordingly, 1 participant failed to be segmented and was excluded. Then we visually checked the images with the mean correlation smaller than three standard deviations from the sample mean, which led to a visual inspection of 4 participants after preprocessing, and subsequently 2 of 4 participants were excluded as there were artifacts observed.

Subsequently, we checked diffusion tensor imaging (DTI) measures using sum of squared error (sse) of diffusion tensor fit that were generated for each participant by FSL DTIFIT. We visually checked the sse images, and excluded 1 participant due to obvious artifacts. This low number is due to previous exclusion during preprocessing.

In the end, there were 344 participants included in our final analyses.

### 4. Statistical analyses using original non-imputed data

#### Case-control difference

Due to 4 individuals lack of full-scale IQ (FIQ), a generalized linear model (GLM) was utilized to examine group difference of the brain's inter-participant variations in linked independent component analysis (LICA) outputs on the 340 participants while regressing out the effect of age, sex, FIQ and scanner site.

We found the same multimodal component (IC58) significant relating to autism (β=-0.189, FDR corrected p=0.040) as using imputed data.

#### Brain-symptom associations

#### Univariate analyses

We used a GLM to explore the univariate associations between each multimodal IC (i.e., no single modality contributed more than 50%) and subscales of Autism Diagnostic Interview-Revise (ADI) (6) and Autism Diagnostic Observational Schedule 2 (ADOS) (7), Social Responsiveness Scale 2nd Edition (SRS) (8), Repetitive Behavior Scale-Revised (RBS) (9), and Short Sensory Profile (SSP) (10) in the autism group correcting for multiple comparison with false discovery rate (FDR) ($p<0.05$).

There are 178 individuals with ADI scores (7 individuals without ADI scores), 179 individuals with ADOS scores (6 individuals missing ADOS scores), 164 individuals with SRS score (21 missing), 156 individuals with RBS score (29 missing) and 107 individuals with SSP score (78 missing) in autism group. Similar to the main findings with imputed data, we did not find any univariate brain-symptom association ($p>0.05$).

#### Multivariate analyses

Subsequently, we utilized canonical correlation analysis (CCA) (11) to detect multivariate association between all multimodal brain independent components (ICs) and all symptom phenotypes in the autism group. The statistical significance of CCA modes was assessed by permutation inference (12). We performed two separate CCA analyses; in the first CCA analyses ($CCA_1$) we included the subscales of ADI and ADOS and in the second ($CCA_2$) we used total scores of parent-rated SRS, RBS, and SSP within autistic individuals. For each CCA's multiple testing correction, we used stepwise cumulative maximum approach, $p<0.05$. The evaluation of the contribution of each IC and each clinical measure to the canonical correlation was according to the structural coefficient of each variable described previously (13).

Due to missing data, there were 153 participants with autism involved in $CCA_1$ (32 participants with autism excluded), and there were 106 participants with autism involved in $CCA_2$ (79 participants with autism excluded). Different from what we found using imputed data (we only find a significant association pattern in $CCA_1$), we separately found a significant multivariate association pattern of $CCA_1$ and $CCA_2$ ($CCA_1$: r=0.827, corrected p=0.003, Figure S1; $CCA_2$: r=0.939, corrected p=0.016, Figure S2). In $CCA_1$, multimodal IC1 (canonical weight: 0.336) and IC5 (canonical weight: 0.065) showed the strong contribution to the correlation with autism core symptoms, and from the symptomatic perspective the ADOS restricted and repetitive behaviors (RRB) subscale showed a high association to the multimodal brain ICs. This multivariate association pattern is similar as the pattern using imputed data, in which IC1 primarily included corpus callosum (CC), bilateral anterior thalamic radiation and corticospinal tract (CST) and IC5 mainly involved shared variation in CC and anterior thalamic radiation. DTI measures dominate in both of these multimodal ICs. And using original data, subscales of ADI have less contributions to the correlation. In $CCA_2$, multimodal IC79 (canonical weight: 0.216) and IC1 (canonical weight: -0.210) contributed most to the correlation, and SSP ranked highest in the three behavior profiles to the association. In IC79, VBM and mode of anisotropy (MO) load most, and gray matter volume variations were seen in bilateral parietal operculum cortex, left precuneus cortex, lateral occipital cortex and a large proportion of cerebellum, while MO variations were mainly found in middle cerebellar peduncle.

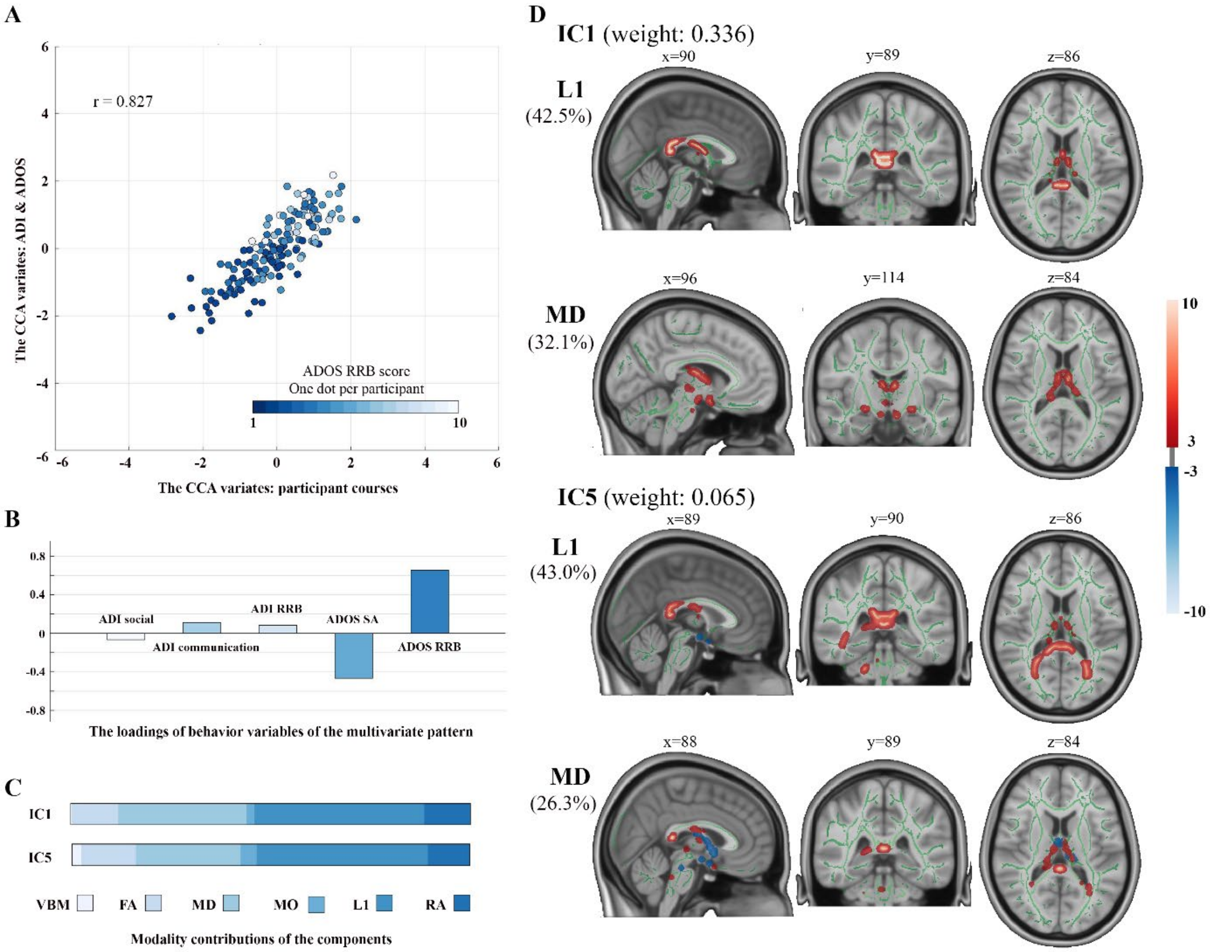

**Figure S1** The multivariate association pattern is found significant between multimodal components and subscales of ADI and ADOS using the original unimputed data. In this pattern, the second contributor of brain components changed to IC5 (instead of IC33 using imputed data, which emphasizes the involvement of the genu and body of corpus callosum). **A** displays the scatterplot of this association pattern, and x, y axes are the pair of variates that derives from CCA. One dot in each participant is coded with gradient color regarding to the RRB subscale of ADOS. **B** demonstrates the loading of each behavior subscale in this association pattern. **C** shows the modality contributions of the multimodal components displayed in **D**. **D** exhibits the two multimodal components contributed most to the association pattern. The weights of each component are showed in the brackets. The spatial maps are thresholded at 3<|Z|<10. All CCAs were only performed in autism group. CCA, canonical correlation analysis; ADI, Autism Diagnostic Interview-

Revised; ADOS, Autism Diagnostic Observational Schedule 2; SA, social affect; RRB, restricted repetitive behavior; IC, independent component; MO, mode of anisotropy; RA, radial diffusivity; MD, mean diffusivity; FA, fractional anisotropy; L1, axial diffusivity; VBM, voxel-based morphometry.

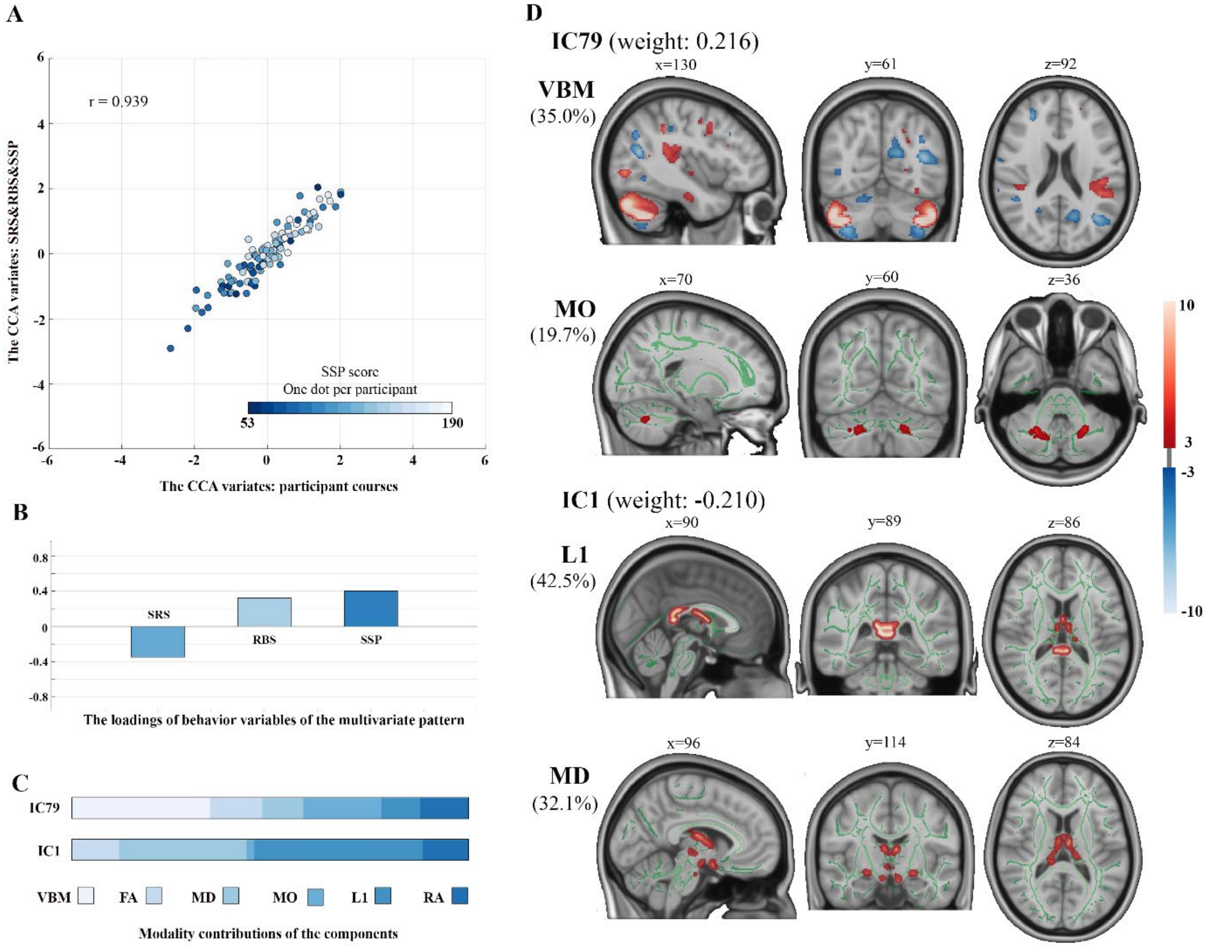

**Figure S2** The multivariate association pattern is found significant between multimodal components and subscales of SRS, RBS and SSP using the original unimputed data (in main text using imputed data, this association did not survive multiple testing correction). **A** displays the scatterplot of this association pattern, and x, y axes are the pair of variates that derives from CCA. One dot in each participant is coded with gradient color regarding to the SSP. **B** demonstrates the loading of each behavior subscale in this association pattern. **C** shows the modality contributions of the multimodal components displayed in **D**. **D** exhibits the two multimodal components contributed most to the association pattern. The weights of each component are showed in the brackets. The spatial maps are thresholded at 3<|Z|<10. All CCAs were only performed in autism group. CCA, canonical correlation analysis; SRS, Social Responsiveness Scale 2nd Edition; RBS, Repetitive Behavior Scale-Revised; SSP, Short Sensory Profile; IC, independent component; MO, mode of anisotropy; RA, radial diffusivity; MD, mean diffusivity; FA, fractional anisotropy; L1, axial diffusivity; VBM, voxel-based morphometry; DTI, diffusion tensor imaging.

### 5. Modality contributions and multimodal index

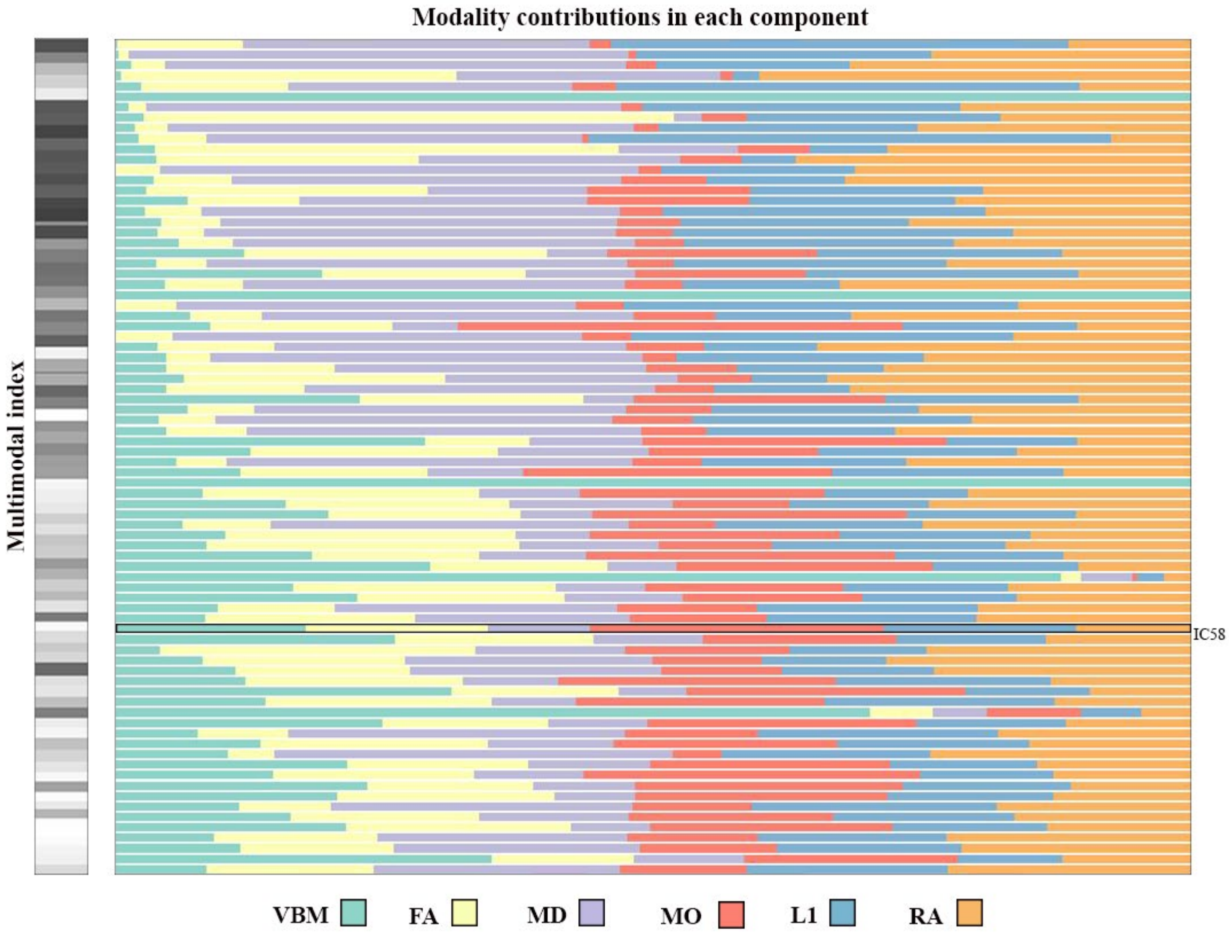

**Figure S3** Modality contributions and multimodal index of each component (80 components, of which there are 75 multimodal components). Multimodal index showing left demonstrates the degree of multimodality, i.e., the closer coded color to dark gray means the more dominant one modality in the components contributes, and in contrast the closer coded color to bright white shows the more equally each modality in the components contributes. MO, mode of anisotropy; RA, radial diffusivity; MD, mean diffusivity; FA, fractional anisotropy; L1, axial diffusivity; VBM, voxel-based morphometry.

### 6. Spatial distribution of VBM and DTI measures in the component with significant group effect

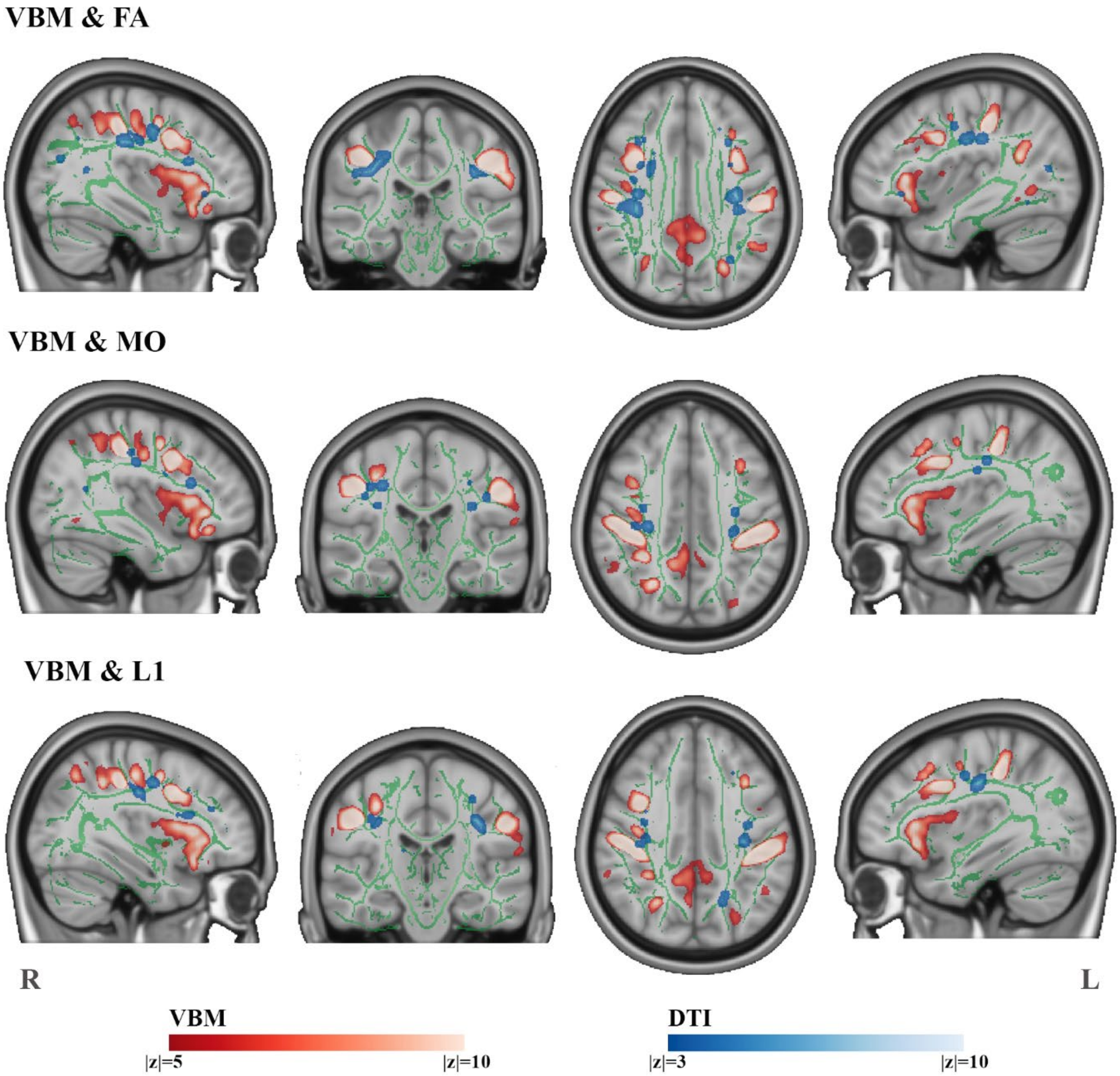

**Figure S4** Spatial distribution of VBM and DTI measures in the component with significant group effect (IC58). The figure displays the top-three loading DTI measures in the component overlaying with the VBM spatial map, which indicates autism-relating variations of gray matter volume and WM tracts that locate around frontal, pre-central and post-central areas are spatially interconnected. The VBM spatial maps are thresholded at 5<|Z|<10, and the DTI spatial maps are thresholded at 3<|Z|<10. VBM, voxel-based morphometry; DTI, diffusion tensor imaging; FA, fractional anisotropy; MO, mode of anisotropy; L1, axial diffusivity; TD, typically developing.

### 7. Scan site effect on the component with significant case-control difference

The participants in our study were recruited in parallel at three collaborating sites (London, Nijmegen and Mannheim). In the main analysis, we included scanner site as a covariate in the GLM. Here, we additionally checked the interaction effect of diagnosis and site to further evaluate the robustness of our autism-related findings. We calculated type-II analysis-of-variance (likelihood-ratio chi-square) for the model including a site-by-diagnosis interaction term (component ~ group*site + age + sex + IQ).

We found that our main effect of diagnosis was robust to the inclusion of site-by-diagnosis in the model ($F=13.110$, p-uncorrected<0.001). There was a modest site-by-diagnosis interaction effect ($F = 6.860$, p-unadjusted=0.032) but no main effect of site ($p>0.300$), suggesting our finding of case-control difference was not significantly driven by site.

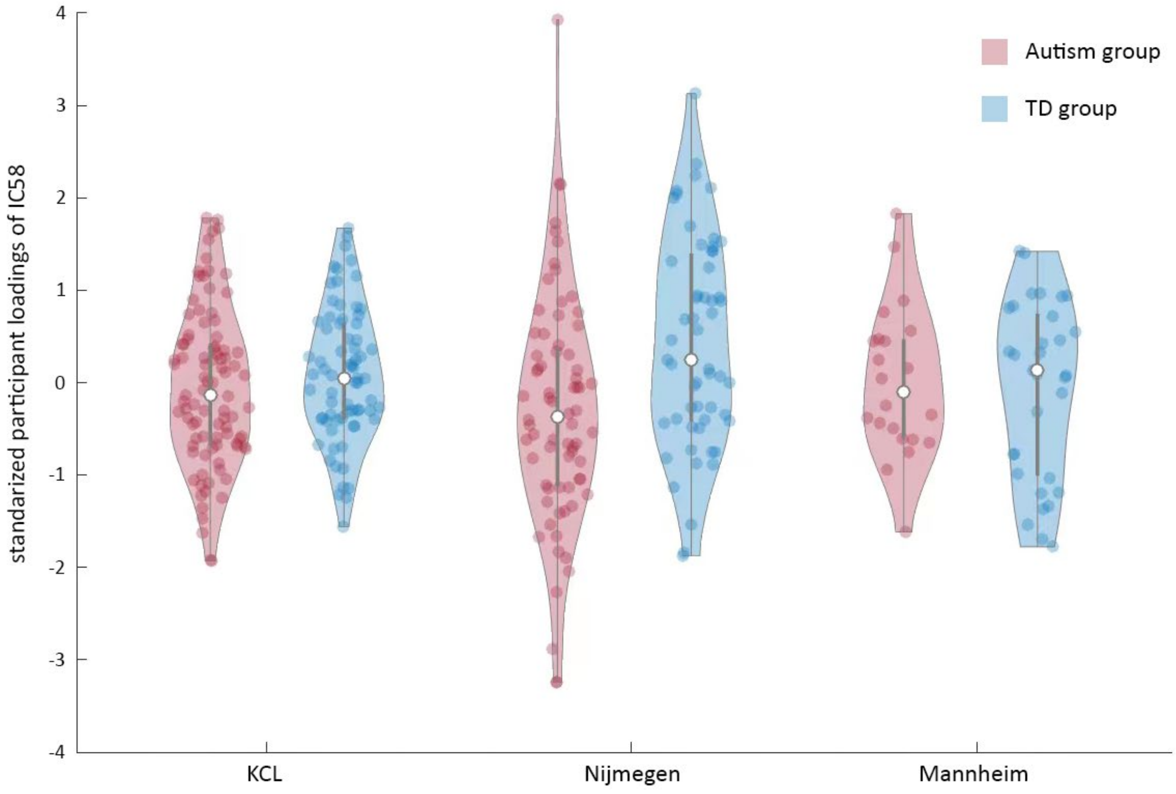

**Figure S5** Distribution of participant loadings of the component with significant group effect (IC58) in the main analyses across scanner sites
